# Independent Continuous Tracking of Multiple Agents in the Human Hippocampus

**DOI:** 10.1101/2025.03.06.641914

**Authors:** Assia Chericoni, Justin M. Fine, Ana G. Chavez, Melissa C. Franch, Elizabeth A. Mickiewicz, Raissa K. Mathura, Joshua Adkinson, Eleonora Bartoli, Joshua Jacobs, Nicole R. Provenza, Andrew J. Watrous, Seng Bum Michael Yoo, Sameer A. Sheth, Benjamin Y. Hayden

## Abstract

The pursuit of fleeing prey is a core element of many species’ behavioral repertoires. It poses the difficult problem of continuous tracking of multiple agents, including both self and others. To understand how this tracking is implemented neurally, we examined responses of hippocampal neurons while humans performed a joystick-controlled continuous prey-pursuit task involving two simultaneously fleeing prey (and, in some cases, a predator) in a virtual open field. We found neural maps encoding the positions of all the agents. All maps were multiplexed in single neurons and were disambiguated by the use of the population coding principle of semi-orthogonal subspaces, which can facilitate cross-agent generalization. Some neurons, more common in the posterior hippocampus, had narrow tuning functions reminiscent of place cells, lower firing rates, and high information per spike; others, which were found in both anterior and posterior hippocampus, had broad tuning functions, higher firing rates, and less information per spike. Semi-orthogonalization was selectively associated with the broadly tuned neurons. These results suggest an answer to the problem of navigational individuation, that is, how mapping codes can distinguish different agents, and establish the neuronavigational foundations of pursuit.

## INTRODUCTION

Pursuit is a foraging behavior involving continuous and interactive navigation with the goal of catching a fleeing prey while avoiding predators (Fabian et al., 2018; Olberg et al., 2000; Stephens & Krebs, 1986; Ydenberg & Dill, 1986). It is an archetypal example of continuous decision-making, in which choice and control are deployed simultaneously (Burge et al., 2025; Cisek and Kalaska, 2010; Gordon et al., 2021; Merel et al., 2015; Yoo et al., 2021A). Moment to moment choices during pursuit require tracking the locations of multiple agents at the same time, including the self, pursued prey, and unpursued prey, and navigating towards interception loci.

Among regions associated with navigation, the hippocampus is the most well studied (Chersi and Burgess, 2015; Ekstrom et al., 2018; Epstein et al., 2017; Kunz et al., 2021; Maguire et al., 2006; Nyberg et al., 2022; O’Keefe and Dostrovsky, 1971; Suthana et al., 2009). The hippocampus contains place cells that track the allocentric position of the self in physical space (Ekstrom et al., 2003; Jacobs et al., 2010; Miller et al., 2013; O’Keefe et al., 1998; Wilson and McNaughton, 1993) and virtual space (Harvey et al., 2009; Mackay et al., 2024). The hippocampus also has a variety of other neurons relevant to pursuit (Behrens et al., 2018). These include neurons that are tuned to positions of external (physical) goals (Brown et al., 2016; Gauthier and Tank, 2018; Kunz et al., 2021; Poucet and Hok, 2017; Watrous et al., 2018), and ‘social’ place cells that track the positions of other agents (Danjo et al., 2018; Forli and Yartsev, 2023; Omer et al., 2018; Rao et al., 2019; Stangl et al., 2021; Zang et al., 2024). The collective existence of these neurons indicates that the hippocampus contains the basic ingredients to track both self and prey in the course of pursuit. However, it does not give insight into how the brain solve the problem of disambiguating representations when faced with multiple distinct agents.

The need to distinguish self from others, and to distinguish multiple others, means that the brain must solve the *individuation problem*. In short, the brain needs a mechanism to know which agent a neuron’s responses refer to. One possibility would involve labelled line coding, in which separate sets of neurons track each single agent. However, such codes tend to be inflexible and have limited capacity for generalization (Barak et al., 2013; Fine et al., 2023; Fusi et al., 2016). In addition to these theoretical concerns, there are empirical ones: most brain areas contain mixed selective codes rather than labeled line codes (Ebitz and Hayden 2021; Fusi et al., 2016; Rigotti et al., 2013; Tye et al., 2024). One way for the brain to handle disambiguation of maps for distinct agents, despite the mixed selectivity, is to represent the maps in semi-orthogonal population subspaces (Elsayed et al., 2016; Johnston et al., 2024; Kaufman et al., 2022; Parthasarathy et al., 2017; Tang et al., 2020; Xie et al., 2022; Yoo and Hayden, 2020). Here, we tested the hypothesis that the hippocampus uses subspace semi-orthogonalization to individuate maps related to distinct agents during continuous pursuit.

We recorded populations of neurons in the hippocampus in thirteen humans performing a virtual pursuit task with two prey and (in a subset of participants) a predator. We found neurons that encode position maps of the self, both prey, and the predator. Some neurons encoded the position of one agent, but the majority were mixed selective for multiple agents; in these mixed selective neurons, maps for different agents were largely unrelated. We found these hippocampal maps can be readily separated into two types: one with narrow spatial extent, reminiscent of place cells, and one with broader less-localized tuning. Narrowly tuned neurons were more prevalent in the posterior hippocampus and encoded position more efficiently, exhibiting greater spatial information per spike despite diminished firing rates, compared to broadly tuned neurons. Notably, the population used subspace semi-orthogonalization for different agents; this coding principle was observed primarily in the broadly tuned neurons. Thus, rather than relying on labeled-line coding, the hippocampus appears to individuate multiple agents through population-level subspace organization, allowing for flexible encoding of distinct yet overlapping spatial representations.

## RESULTS

### Prey pursuit in humans

Human participants (n=13) performed the *prey-pursuit task* (**Figure 1A**, **Methods;** Yoo et al., 2020). On each trial, the participant used a joystick to continuously move the position of an avatar (yellow circle) in a rectangular field displayed on a computer screen (**<u>Supplementary</u> <u>Video</u>, Figure 1A** and **B**). The participant had up to 20 seconds to capture fleeing prey (colored squares) to obtain points. Prey avoided the avatar with a deterministic strategy that combined repulsion from the avatar’s current position with repulsion from the walls of the field (**Methods**). The prey items were drawn randomly on each trial from a set of three that differed in maximum velocity and reward size.

**Figure 1.**
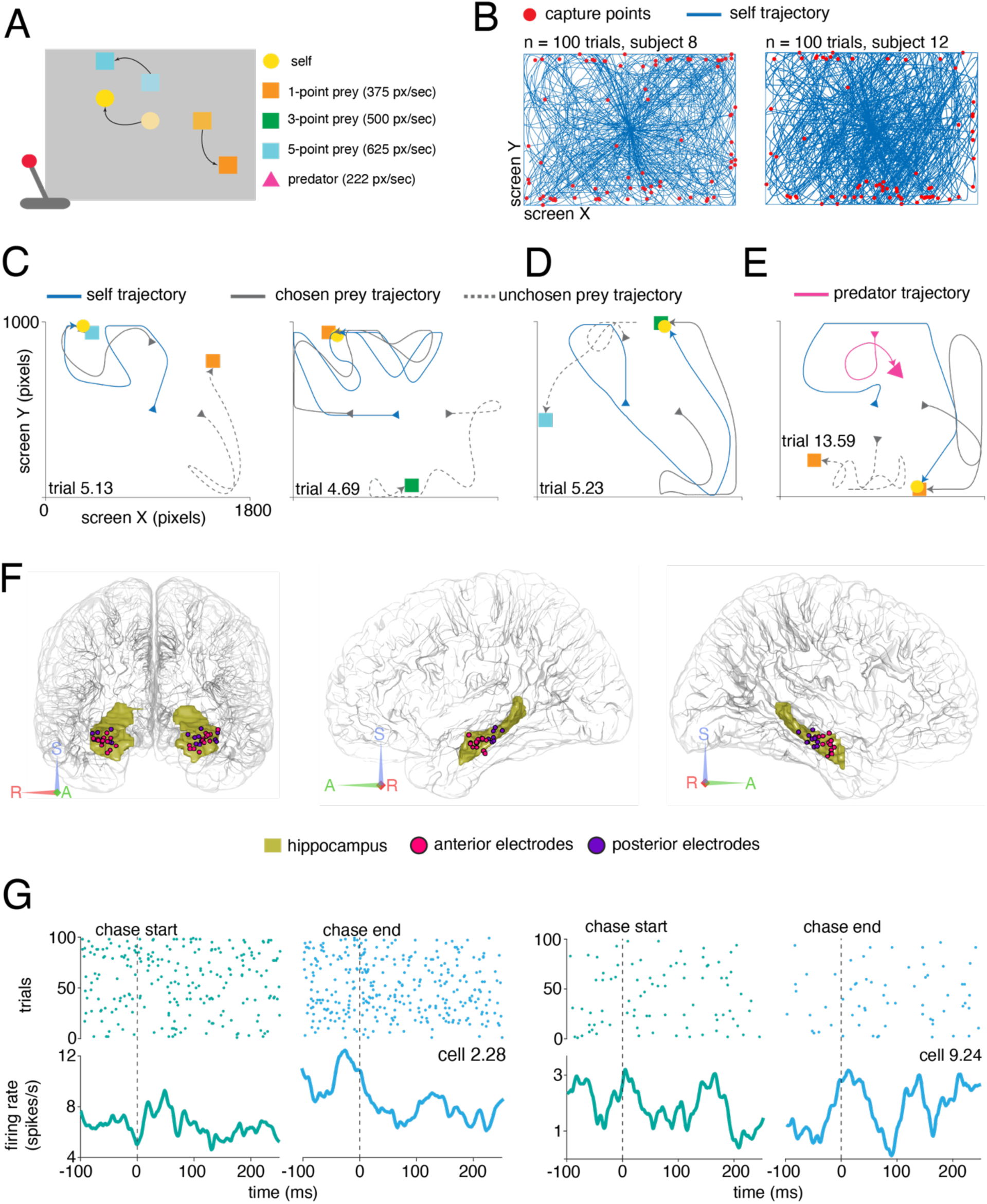
Task design and neural recordings. **A**, Schematic of the prey-pursuit task. Participants use a joystick to control the position of an avatar (yellow circle) on a computer screen to capture prey (squares) and score points. For some participants, we also included predators (**Methods**). **B,** Example sessions from two participants; blue lines indicate participants’ trajectories on each trial overlaid; red dots indicate points of prey capture. **C, D, E,** Typical example trials from different participants. Trials are identified as *participant number.trial number*. Continuous gray lines indicate the chosen prey trajectories and dashed gray lines the unchosen prey trajectories. Predators’ trajectories are reported in magenta. **D,** Example trial in which the participant switched from pursuing one prey to another. **E,** Example of trial with predator - capture from the predator leads to point loss. **F,** Recording sites of hippocampal neurons from all 13 participants. Recording sites within the anterior hippocampus are reported in magenta while recording sites within the posterior hippocampus in purple. **G,** Peri-stimulus time histograms and raster plots demonstrating responses to chase start and chase end from two hippocampal neurons from participant 2 (left column) and participant 9 (right column). Dashed vertical lines represent the beginning and the end of the chase. Plots display mean firing rates.

Each trial began with one or two prey appearing at one of the cardinal points (**Figure 1A**). In two-prey trials (81% of trials), the participant was free to decide which prey to pursue at any moment (**Figure 1C, D**). Participants successfully captured the prey in 73.72% of trials and, on successful trials, did so in an average of 7.51 seconds (variance: 1.67 seconds) with an average reaction time of 0.87 seconds (variance: 0.01 seconds), measured as the interval between the agents’ appearance on the screen and the participant’s first move. Participants’ performance did not depend significantly on prey type (trial length x reward level, p = 0.16; reaction time x reward level, p = 0.51). In three participants, we employed a variant of the task in which, in addition to the prey, there were also predators pursuing the participant’s avatar (**Figure 1E**). In this case, the predator used a simple distance-minimizing pursuit strategy. Our participants successfully evaded capture by the predator in 91.67% of trials.

We recorded responses of 390 neurons in the hippocampus while participants performed this task (average n=30 neurons per participant). Of these neurons, 96 were also recorded in the variant of the task with a predator. Of all 390 neurons, roughly half (n=199) were in anterior hippocampus and the remainder (n=191) were in posterior hippocampus (**Figure 1F**). We defined the border between these regions as a coronal plane along the longitudinal hippocampal axis (y = -20, MNI). This border is largely consistent with that used in previous studies (e.g., Poppenk et al., 2013). Average firing rates aligned to trial stop and start show some intriguing patterns (**Figure 1G**). Overall, however, neurons had complex selectivities that were not readily explainable in terms of trial start and stop. We therefore examined responses as a function of location of each agent in the virtual space of the task.

### Hippocampal maps for positions of self, prey, and predators

To estimate mapping functions in these neurons, we used the Poisson generalized linear model procedure developed by Hardcastle et al. (2017). This approach fits tuning models to neuronal responses without any a priori assumptions about the shape of the tuning surface. For this analysis, and all the subsequent ones, we concatenated all the successful trials involving two prey. (Single prey trials showed similar results and are not described here). To describe our results here, we use the term *chosen prey* for the prey that was ultimately captured, while the other was the *unchosen prey* (**Figure 1C, D**).

We found 37.7% (n=147/390) of neurons map the position of the self, while 33.9% (n=132/390) map the position of the chosen prey, and 24.4% (n=95/390) map the position of the unchosen prey (**Figure 2A, B, D, F, G**). In the neurons that we recorded during predator trials, 27.08% (n=26/96) map the position of the predator (**Figure 2E**). We found that 15.1% (n=59/390) of the neurons encode the position of any two agents, and 14.4% (n=56/390) map all three agents (**Figure 2B**). Similarly, during predator trials, 15.6% of the neurons (n=15/96) map the position of two or three agents, with 8.3% (8/96) mapping the position of all four agents. These proportions are all higher than would be expected by chance (p<0.001, binomial test).

**Figure 2.**
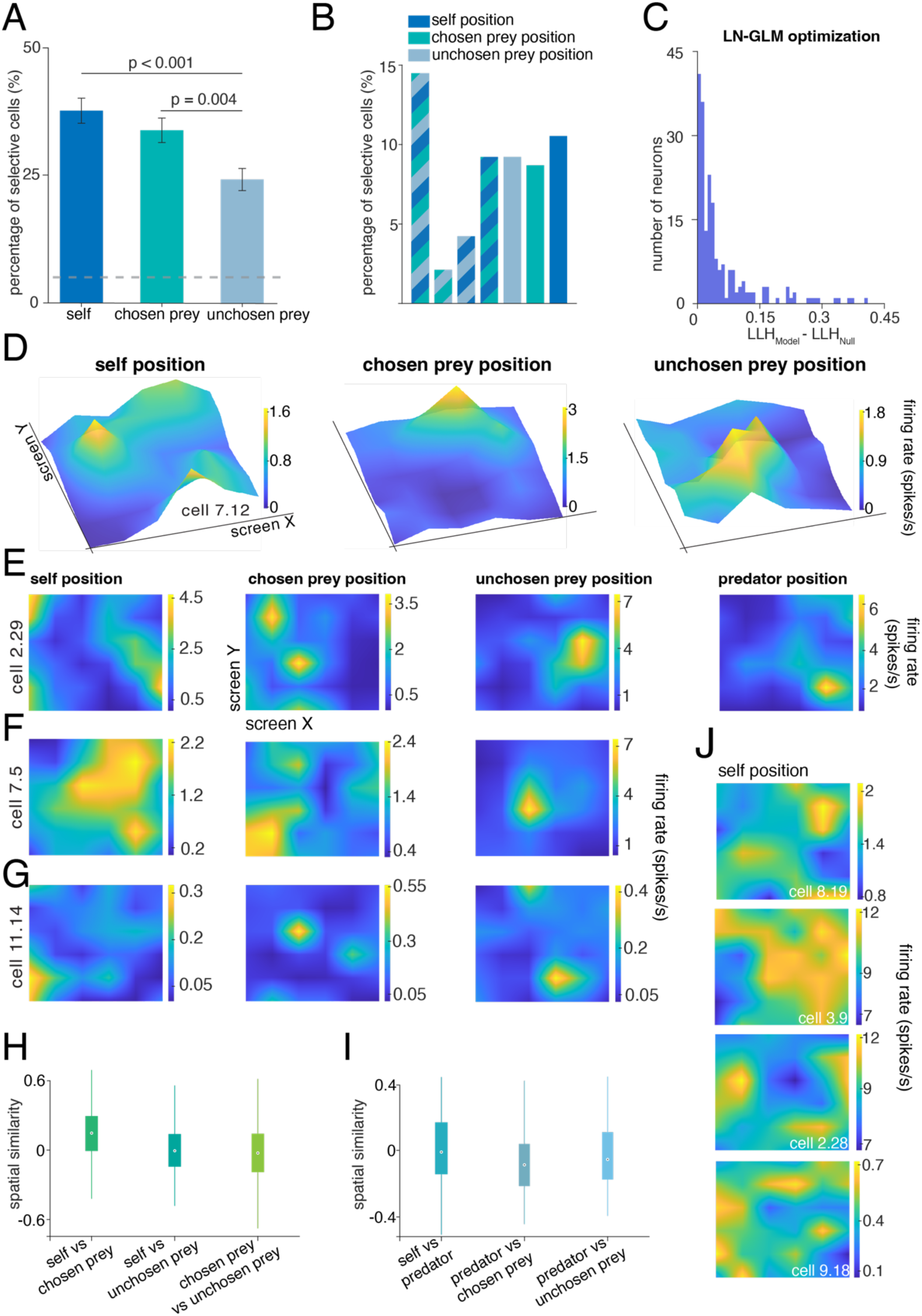
Hippocampal mapping functions for self and prey. **A**, Bar plot showing the percentage of neurons tuned to agent positions according to the LN-GLM approach. In total 51.79% (202/390) of the neurons were selective for any of the agents’ positions. Proportions do not sum up to hundred as one neuron may be tuned to more than one agent’s position. Dashed horizontal grey line represents chance level, which is 5%. Error bars represent standard errors. Significance was determined using a chi-square test for proportions, with a significance threshold of 0.05. **B,** Barplot showing the proportion of neurons tuned to more than one agent’s position. For instance, the first bar from the left shows that 14.36% of the neurons were selective for the position of all three agents. **C,** Quantification of model performance using log-likelihood (LLH) increase. The histogram shows LLH increase values for neurons significantly tuned to at least one agent’s position. LLH increase was computed as the improvement over a null model assuming a constant firing rate, using 10-fold cross-validation. Only neurons where the best-fit model significantly outperformed the null model (Wilcoxon signed-rank test, p < 0.05) are shown. LLH increase values were normalized by spike count and converted to bits per spike using log base 2 scaling. **D, E, F, G,** Three-dimensional (top row) and two-dimensional (bottom rows) representations of spatial firing rate maps for neurons significantly tuned to the position of all the agents. Each row corresponds to a single neuron, while each column represents the firing activity relative to different agents (self, chosen prey, unchosen prey, and predator when present). Yellower regions indicate locations where the neuron exhibited higher firing rates.

Next, we asked whether hippocampal neurons have different maps for the different agents. To quantify the relationship between maps, we used a *spatial similarity index* (SPAEF, Koch et al., 2018, **Methods**), which measures the correlation between spatial representations. A value of zero indicates full orthogonality between maps, while values closer to +1 or -1, indicate correlation or anti-correlation between maps, respectively.

The mean spatial similarity between self and chosen prey maps is 0.14. This value is very low, but is nonetheless greater than zero (p = 0.002, **Figure 2H**, **Methods**) and below noise ceiling (p < 0.001). Thus, these maps appear to be largely, but not entirely distinct. The mean spatial similarity between self and unchosen prey maps is even lower, but is still different from zero (SPAEF = 0.002, p < 0.001, **Figure 2H**). The chosen and unchosen prey maps are weakly, but significantly anti-correlated (SPAEF = -0.02, p = 0.001, **Figure 2H**). Also in this case, both values were below noise ceiling (p < 0.001). Finally, in the subset of neurons in which we had predator data, we found that the spatial similarity between self and predator was again slightly positive but not different from zero (SPAEF = 0.006, p = 0.79, **Figure 2I**). These single-neuron SPAEF results are reminiscent of the idea of semi-collinearity, in which coding at the population level shows a mixture of orthogonality and collinearity (Johnston et al., 2024). We therefore next tested this population level idea directly.

Data points were interpolated for visualization purposes only. **H,** Boxplots representing the median spatial similarity between maps across different agent representations, errorbars represent the standard error. Spatial similarity values near zero indicate orthogonality (low similarity) between maps. All the neurons are shown. **J,** Example maps from neurons that were significantly tuned to self position.

### Narrow and broad tuning curves for spatial position

We next surveyed the distribution of tuning curve shapes. Some hippocampal neurons have selectivity for positions that are localized to a specific location, akin to the narrow maps of place cells (**Figure 2D, E, G**). However, others have broader response functions that are not as narrowly localized, but that nonetheless carry strong spatial information (**Figure 2F, J**).

We performed a k-means clustering of self-position tuning functions (**Methods**). We validated the clustering output using silhouette scores, which can provide an estimate of the most likely number of true clusters (Rousseeuw, 1987). This analysis shows that only two clusters are needed (mean silhouette value = 0.6, **Figure 3A, B**). In our population of neurons, 70.5% (n=275/390) belong to cluster 1 (C1) which has neurons with broad spatial tuning; the remaining 29.5% (n=115/390) are in cluster 2 (C2), which has neurons with narrow place cell-like tuning (**Figure 3A, B**).

**Figure 3.**
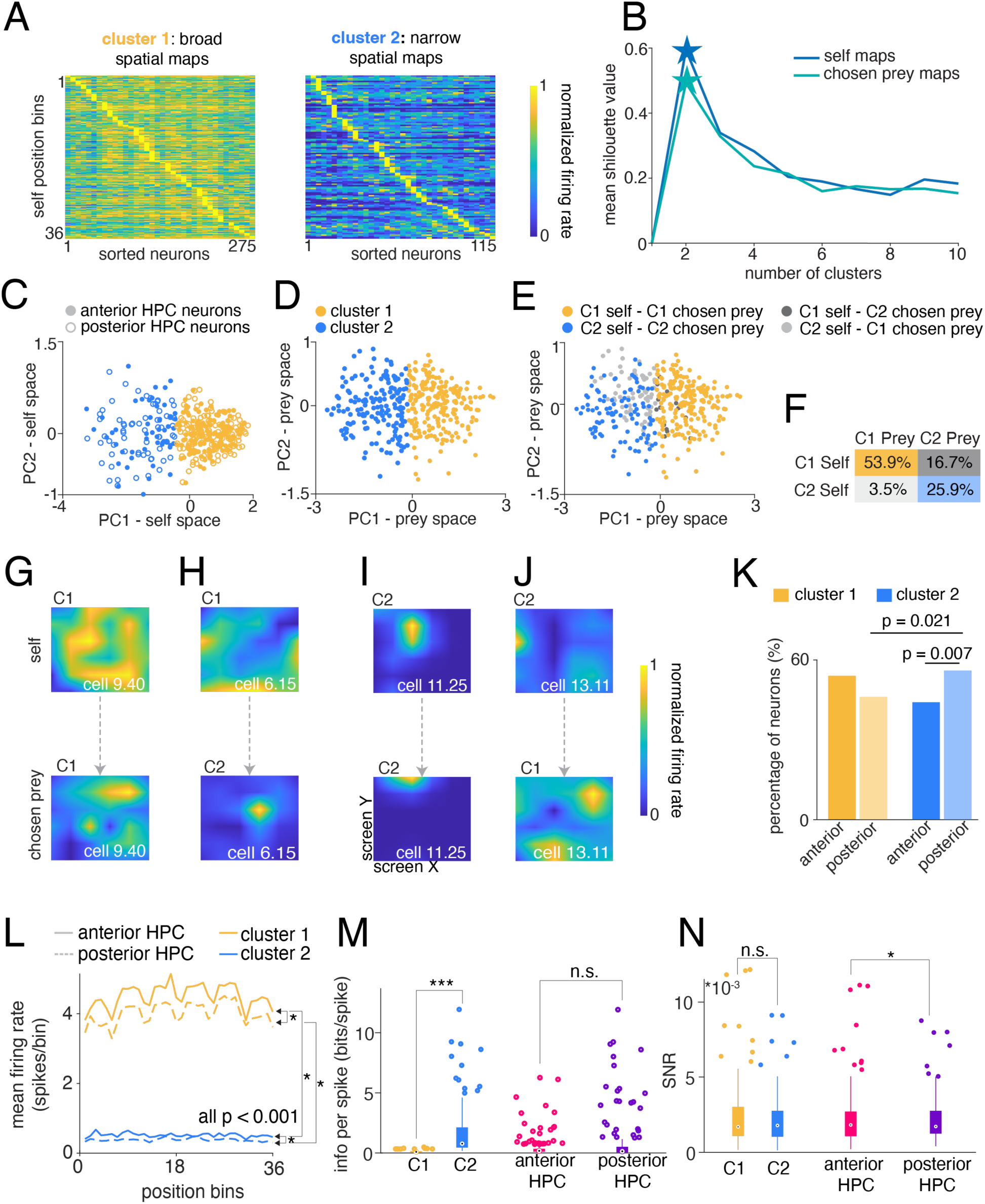
Hippocampal tuning profiles are heterogeneous. **A**, Heatmaps depicting the two clusters of neurons identified using k-means clustering. Each matrix represents unfolded neuron maps, where rows correspond to position bins and columns to individual neurons. Neurons are sorted according to their peak firing positions (bins). Firing rates are normalized within neurons between 0 and 1. **B,** Silhouette scores used to assess clustering quality and determine the optimal number of clusters for both self and chosen prey maps. Stars indicate the selected number of clusters. **C,** Neurons projected onto the PC space derived from self-position maps, colored according to k-means clustering performed on self maps. **D,** Same as in C, but PCA and k-means were performed on chosen prey maps. **E,** Neurons projected onto the chosen prey PCs space, now colored by cross-referencing k-means clustering results from self and chosen prey maps. **F,** Proportion of neurons assigned to each cluster based on their self and chosen prey representation. The table highlights the percentage of neurons that remained in the same cluster across self and chosen prey representations, as well as those that switched clusters between conditions. **G, H, I, J,** Example neurons demonstrating shared (G, I) or distinct (H, J) spatial representations for self and chosen prey. Each column corresponds to a single neuron, with the top rows showing spatial tuning to self and the bottom row tuning to chosen prey. **K,** Comparison of neuron distribution between anterior and posterior hippocampus across clusters. Significance is based on one sample z-test for proportions. **L,** Plot representing the mean firing rate across a 6x6 grid (36 bins - x axis). All p < 0.001, Wilcoxon rank sum test. **M,** Boxplot representing the spatial information per spike computed as the mutual information between the spatial location and the spike train, p < 0.001, Wilcoxon rank sum test. **N,** Boxplot representing the SNR computed as the ratio between the variance of the tuning and the average firing rate, where a higher SNR means a neuron fires more reliably in specific locations; p = 0.02, Wilcoxon rank sum test.

We applied principal component analysis (PCA) on the self-position tuning functions and visualized the population distribution in a lower dimensional space (**Methods**). Neurons belonging to cluster 1 and cluster 2 are clearly separable along the first principal component (PC1), which represents the overall *spatial extent* of the tuning, while the second principal component (PC2) may capture differences in neurons’ preferred firing locations, signifying *spatial preference* (**Figure 3C**). Indeed, neurons with broad spatial representations are distributed on the right side of PC1, with little variability along both PC1 and PC2, which may indicate greater spatial extent and lower spatial preference. That is, firing patterns are more widespread and heterogeneous across multiple spatial locations (**Figure 3C**). In contrast, neurons with narrow spatial representations were more dispersed along both PC1 and PC2, reflecting greater variability in their tuning peaks - spatial extent - and spatial preference (**Figure 3C**).

We performed the same clustering procedure, but this time using chosen prey’s position tuning functions (**Methods**). As with self-position, the silhouette test showed optimal results with k = 2 (mean silhouette value = 0.5, **Figure 3B**), confirming that population response relies on broad and narrow maps to represent position (**Figure 3D**). For these chosen prey maps, we found that 57.44% (n=224/390) belong to cluster 1, while the remaining 42.56% (n=166/390) belong to cluster 2. The overwhelming majority of neurons (79.80%) consistently fall into the same cluster for self- and prey-position (**Figure 3E, F, G, I**), whereas the remaining 20.20% of neurons switch between broad and narrow spatial maps (**Figure 3E, F, H, J**). Repeating the same analysis on both self and chosen prey maps, first applying PCA followed by k-means clustering, yielded similar results.

We found that broad and narrow neuron maps are differentially distributed along the longitudinal axis of the hippocampus. Specifically, we observed a greater preponderance of broadly tuned neurons in the anterior hippocampus and a greater preponderance of neurons with narrow spatial tuning in the posterior hippocampus (**Figure 3K**). Among cluster 1 neurons, 53.80% localized rostrally with the remaining 46.20% caudally, with no significance difference between them (one sample z-test for proportions: p = 0.99). Among cluster 2 neurons, 44.30% are found rostrally and 55.70% caudally. While this difference is modest, it is statistically significant (one sample z-test for proportions: p = 0.007; **Figure 3K**). This result remained unchanged even when considering only the significantly tuned neurons. Out of the cluster 2 tuned neurons, 42.10% localize in the anterior hippocampus and 57.90% in the posterior (one sample z-test for proportions: p = 0.003; data not shown). Moreover, within the posterior hippocampus, we found a higher proportion of cluster 2 neurons compared to cluster 1 (one sample z-test for proportions: p = 0.021; **Figure 3K**). Within the anterior hippocampus, instead, there was no significant difference between the proportion of cluster 1 and cluster 2 neurons (one sample z-test for proportions: p = 0.98, **Figure 3K**). Together, these results confirm theories that anterior and posterior hippocampus have measurable functional differences.

Last, we examined how these two distinct categories of neurons fire across space along with the associated signal-to-noise ratio (SNR), defined as the tuning curve variance normalized by firing rate (**Methods**). We found that neurons carrying broad spatial representation exhibit higher firing rates but carry less spatial information compared to narrowly tuned neurons (ranksum, p < 0.001; **Figure 3L, M**). Nonetheless, the SNR remained comparable across clusters (**Figure 3N**), indicating that the firing rate and information per spike roughly cancelled each other out. However, we did find that neurons in the posterior hippocampus had lower firing rates but higher SNR compared to anterior hippocampal neurons (**Figure 3L, N**). These findings highlight distinct encoding strategies along the longitudinal hippocampal axis.

### Hippocampal neurons use semi-orthogonal maps to disambiguate agents

Theoretical and empirical data support the idea that neural populations can use orthogonal neural subspaces to partition information encoded in the same neurons (Ebitz and Hayden, 2021; Elsayed et al., 2016; Kaufman et al., 2022; Panichello and Buschman, 2021; Tang et al., 2020; Xie et al., 2022; Yoo and Hayden, 2020). Moreover, blending orthogonal and collinear subspaces, producing semi-orthogonal subspaces, allows for discrimination while enabling cross-categorical generalization (**Figure 4A-D**; Barak et al., 2013; Bernardi et al., 2020; Johnston et al., 2024).

**Figure 4.**
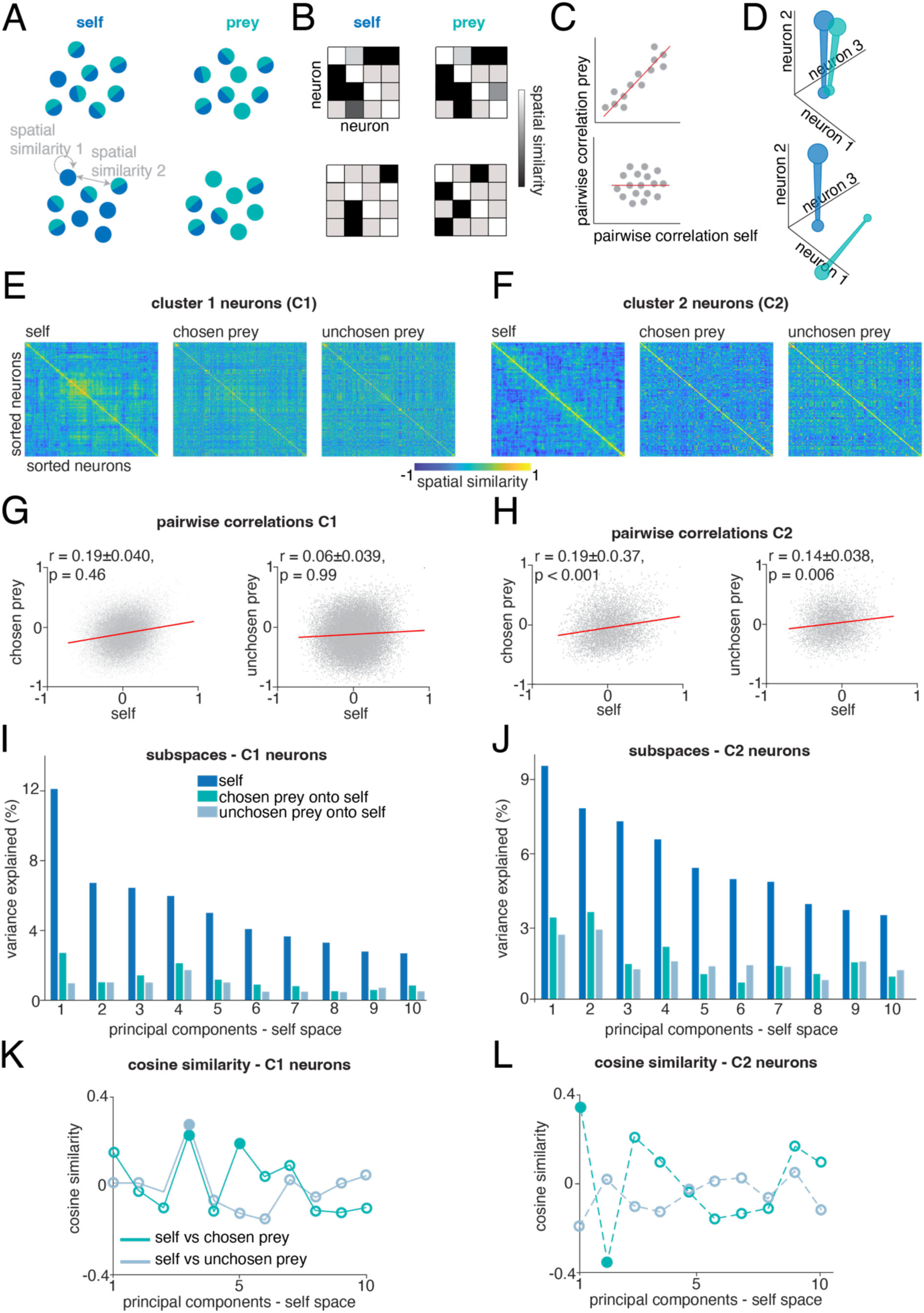
Neural subspaces allow for agent generalization. **A, B, C, D**, Schematic of neural subspaces orthogonalization. **A,** Cartoon showing activity of ensemble neurons. Each neuron maps the position of one or more agents. **B, C,** Spatial similarity matrix and pairwise correlation; uncorrelated activity results in subspace orthogonalization (bottom row) and vice versa (upper row). **D,** Blue represents the self subspace, while green represents the prey subspace. Correlated neural activity results in collinear subspaces (top row) and viceversa (bottom row). **E,** Spatial similarity matrices for all the possible neuron pairs in cluster 1 for self and prey. Neurons of each matrix are sorted using a hierarchical clustering algorithm applied on the self matrix only. Z axis indicates the strength of correlation between the overall average activities of neurons pairs. Lack of obvious structure in the self matrix reflects strong functional reorganization between self and prey. **F,** Same as in E, but for cluster 2 neurons. **G,** Pairwise correlation for each pair of neurons in cluster 1 for self and prey. The lack of correlation indicates that the relationship between neurons changes across agents. Reported values indicate Pearson’s correlation coefficient with 95% confidence intervals, and permutation-based p-values were computed using a two-tailed test, with significance considered at p<0.05. **H,** Same as in E, but significant correlations indicate that the relationship between neuron pairs in cluster 2 is preserved across agents, suggesting a shared representational structure. **I,** Percent variance explained by each of self-PCs for prey maps projected onto the self. The low height of the green and grey bars relative to the blue ones illustrates the poor match between self and prey subspaces. **J,** As in I, but for cluster 2 neurons. **K,** Cosine similarity between self and prey PCs, values closer to zero indicate orthogonality between PCs. Filled dots represent nonzero values (two-tailed test, with significance considered at p<0.05). **L,** as in K, but for cluster 2 neurons.

To test the hypothesis that hippocampal neural populations use semi-orthogonal subspaces to disentangle agents’ maps, we adapted a method previously developed to study motor regions (Elsayed et al., 2016). This analysis quantifies the degree of overlap between spatial representations for self and prey, by projecting the population activity onto low-dimensional subspaces that capture most of the variance between agents’ representations (**Methods**). We applied this analysis separately to both clusters. We found that neurons with broad spatial maps (cluster 1) use distinct, semi-orthogonal subspaces to individuate agents’ locations. We also found that neurons with narrow spatial maps (cluster 2) represent self and prey on nearly collinear subspaces.

In more detail, we started by analyzing the correlation structure for self and prey representations. For each agent, we computed the spatial similarity between the tuning surfaces for each pair of neurons (**Figure 4E, F**). Each entry in the resulting *spatial similarity matrix* represents the degree of spatial similarity between the response patterns of two neurons. For C1 neurons, the matrix structure changes markedly across agents (**Figure 4E**), whereas for C2 neurons, it remains largely consistent (**Figure 4F**). Indeed, when we compared the entries of the self matrix against the entries of the prey matrix, we found little or no correlation (C1: not different from chance, determined by pseudocorrelation, self vs chosen prey: R2 = 0.19 ± 0.040, p = 0.46; self vs unchosen prey: R2 = 0.06 ± 0.039, p = 0.99; two-sided permutation test, α = 0.05; **Figure 4G**; C2: self vs chosen prey: R² = 0.19 ± 0.037, p < 0.001; self vs unchosen prey: R² = 0.14 ± 0.038, p = 0.006, **Figure 4H**). Thus, for a pair of neurons with broad spatial maps (cluster 1), the similarity of their responses to self-locations does not inform the similarity of their responses to prey locations. A possibility is that these neurons individuate agents in space with some dimensions dedicated to self-only, some to prey-only, and some to both - aligning with the hypothesis of semi-orthogonality between neural subspaces.

To understand if these subspaces were semi-orthogonal, we applied eigendecomposition to the self spatial similarity matrix and individuated the top self-principal components (PCs). If self and prey maps are semi-orthogonal, the self-PCs should capture very little prey variance (**Methods**). To assess if self-PCs can significantly capture the prey maps variance, we computed variance explain noise ceiling and confidence intervals through half-split cross-validation (CV) (**Methods**). For each split, we used half of the self spatial maps (train half) to compute the self-PCs. We then use these self-PCs to project the test half’s spatial maps for both self (within-self variance) and prey (cross-agents variance). The distribution of within-self variances gives an estimate of noise ceiling, while the cross-agents variance explained quantifies how well self-PCs capture prey maps variance. The variability of both within-self and across-agents variance provides confidence intervals for each agent’s projections.

For cluster 1 neurons, we found that the prey variance explained was significantly lower than noise ceiling (p < 0.001, for PC1 to PC10), though confidence intervals did not overlap between self and prey projections for PC1 only. That is, self-PC1 carries information unique to the self maps. Overlapping confidence intervals for the remaining components implied that self and prey representations share non-zero variance (**Figure 4I**).

Lastly, to estimate the degree of orthogonality between agents’ neural subspaces, we identified the top prey-PCs, and estimated the cosine similarity between self and prey -PCs. A value of cosine similarity close to zero means orthogonality, while values closer to 1 or -1 collinearity. The cosine similarity values between the self-PCs and prey-PCs, fluctuate around zero, confirming weak alignment between PCs, i.e., semi-orthogonality between subspaces (**Figure 4K**). In summary, these results tell us that neurons with broad spatial tuning individuate prey and self using semi-orthogonal neural subspaces.

The same decomposition analysis performed on neurons with sparse spatial tuning (cluster 2), yielded slightly different results. Given the significant correlations between self and prey representations, a potential explanation is that agents lie on collinear subspaces. We found that also in this case, the prey variance explained is significantly lower than noise ceiling (p < 0.001, for PC1 to PC10), but confidence intervals overlap across all the PCs, indicating shared variance between self and prey maps (**Figure 4J**). The cosine similarity between the top two self-PCs and chosen prey-PCs significantly differs from zero (PCs1: 0.78, p = 0.02; PCs2: -0.36, p = 0.04, **Figure 4L**). Our results indicate that neurons with narrow spatial maps represent self and prey in nearly collinear subspaces, and that subspace semi-orthogonalization in hippocampal neurons is primarily driven by neural populations with broad spatial tuning.

## DISCUSSION

Here we find, in a virtual prey pursuit task, that neurons in the human hippocampus can track positions of multiple agents (self, two prey, and a predator) simultaneously using distinct maps. Rather than use a labelled line code (in which a given neuron’s responses correspond to the position of a single agent), maps are multiplexed. This multiplexing of information raises an *individuation problem*: if firing rate of a neuron changes, which of the tracked agents does that change refer to? Our data suggest that the hippocampus solves this problem by recourse to *semi-orthogonal subspaces*, which blend orthogonal and collinear subspaces. Subspace partition allows for simultaneous separation of information (thus solving the individuation problem) while also allowing for cross-agent generalization, greatly facilitating flexible learning (Barak et al., 2013; Fusi et al., 2016; Johnston et al., 2024). These results, then, build on recent findings showing that rat and bat hippocampus contains cells whose firing rates encode the positions of conspecifics in the environment. Specifically, we find that cells with similar functional properties can be found in the human hippocampus, and moreover, that they can flexibly partition information to achieve individuation.

We find evidence for two categories of cells, based on the shape of their tuning surface for location (broad and narrow). The cells in the smaller cluster 2 have narrow highly localized regions of strong activity, which give them a clear featural resemblance to place cells. However, the cells in the larger cluster 1 have a distinct lack of localized tuning, but nonetheless still carry information about spatial position. These cells have some resemblance to non-grid cells in the rat entorhinal cortex, which were discovered using the same mapping algorithm we employ here (Hardcastle et al., 2017 see also Diehl et al., 2017). Moreover, the category a cell falls in for self-maps is generally the same category as it falls in for other maps, suggesting map shape may be an intrinsic property of neurons, rather than an arbitrary feature. Surprisingly, the subspace semi-orthogonalization appears mostly to be a property of the cluster 1 neurons, suggesting that an overly strict focus on place-cells with focal tuning may result in ignoring important functional features of hippocampal maps.

Our results show a modest but highly significant difference in the distribution of functional cell types in the anterior (fewer cluster 2) and posterior (more cluster 2) hippocampus. These results relate to ongoing discussions about the potential functional differences between these regions (which are homologous to the rodent ventral and dorsal hippocampi, respectively, Strange et al., 2014; To et al., 2024). The increased number of place-cell like coding in posterior hippocampus, then, recapitulates known findings from rats (e.g., Jung et al., 1994; Kjelstrup et al., 2018). Finally, the localization of partitioning to the more anterior cluster 1 neurons aligns this part of the hippocampus with the kind of flexible adaptive cognition associated with prefrontal cortex, with which anterior hippocampus has strong connections (Dalton et al., 2022). Nonetheless, the differences we observe are modest, and, overall, our results endorse functional continuity of the human hippocampus.

Our results relate to a previous series of three papers from our laboratory, in which macaques performed a similar task (Yoo et al., 2020 and 2021B; Fine et al., 2024). In those papers, we found that dorsal anterior cingulate cortex (dACC) actively predicts the future position of pursued prey about 700 ms into the future (Yoo et al., 2020), also tracks a suite of allocentric and egocentric variables related to the task (Yoo et al., 2021B), and corresponds with inferred decision variables in the task (Fine et al., 2024). Relative to macaques, human behavior is quite different - most critically, our macaque subjects needed at least three months to learn the task, whereas humans need no training. Moreover, humans show evidence of highly efficient behavioral strategies, such as cornering the prey, that macaques do not seem to learn, even with months of training. In any case, our results here, focusing on the individuation problem, which was not addressed in our earlier work, complement and extend our earlier findings.

We find it interesting that hippocampus actively tracks the position of the unpursued prey. In our task, subjects typically select a single prey and pursue it for several seconds until capture, leaving the other, unchosen one, ostensibly irrelevant. Nonetheless, from the perspective of the hippocampus, this unselected prey shows only modest evidence of deprioritization. (Namely, hippocampus still tracks the unpursued prey, although less robustly as it tracks the pursued prey). The goal of the task is not, however, simply to track the preferred prey; instead the player is allowed to switch to following the alternative (and sometimes does so, when, for example, the unpursued prey comes nearby). Thus, from the perspective of choice, although not behavior, monitoring the position of the unpursued prey is crucial: because switching is always a possibility, continuing to focus on the pursued prey represents a decision to not switch. Our results are reminiscent of theories in which the hippocampus maps potential alternatives to current goals, rather than being limited to representing only the goals themselves.

## Acknowledgements

We thank George Kokalas and Victoria Gates for invaluable assistance.

## METHODS

### Human intracranial neurophysiology

Experimental data were recorded from 13 adult patients (6 males and 7 females) undergoing intracranial monitoring for epilepsy. The hippocampus was not a seizure focus area of any patients included in the study. Single neuron data were recorded from stereotactic (sEEG) probes, specifically AdTech Medical probes in a Behnke-Fried configuration. Each patient had an average of 3 probes terminating in left and right hippocampus. Electrode locations are verified by co-registered pre-operative MRI and post-operative CT scans. Each probe includes 8 microwires, each with 8 contacts, specifically designed for recording single-neuron activity. Single neuron data were recorded using a 512-channel Blackrock Microsystems Neuroport system sampled at 30 kHz. To identify single neuron action potentials, the raw traces were spike-sorted using the WaveClus sorting algorithm (Chaure et al., 2018) ^and^ then manually evaluated. Noise was removed and each signal was classified as multi or single unit using several criteria: consistent spike waveforms, waveform shape (slope, amplitude, trough-to-peak), and exponentially decaying ISI histogram with no ISI shorter than the refractory period (1 ms). The analyses here used only single unit activity.

### Electrode visualization

Electrodes were localized using the software pipeline intracranial Electrode Visualization (iELVis, Groppe et al., 2017) and plotted across patients on an average brain using Reproducible Analysis & Visualization of iEEG (RAVE, Magnotti et al., 2020). For each patient, DICOM images of the preoperative T1 anatomical MRI and the postoperative Stealth CT scans were acquired and converted to NIfTI format (Li et al., 2016). The CT was aligned to MRI space using FSL (Jenkinson and Smith, 2001; Jenkinson et al, 2002). The resulting coregistered CT was loaded into BioImage Suite (version 3.5β1) (Joshi et al., 2001) and the electrode contacts were manually localized. Electrodes coordinates were converted to patient native space using iELVis MATLAB functions (Joshi et al., 2011; Yang et al., 2012) and plotted on the Freesurfer (version 7.4.1, Dale et al., 1999) reconstructed brain surface. Microelectrode coordinates are taken from the first (deepest) macro contact on the Ad-Tech Behnke Fried depth electrodes. RAVE was used to transform each patient’s brain and electrode coordinates into MNI152 average space (Magnotti et al., 2020). The average coordinates were plotted together on a glass brain with the hippocampus segmentation and colored by location within the hippocampus.

### The prey-pursuit task

The task used here is similar but not identical to one we have previously used in macaques (Yoo et al., 2020A and 2021; Fine et al., 2024; **<u>Supplementary Video</u>**). At the beginning of each trial, two shapes appeared on a gray background (RGB: 128/128/128) on a standard computer monitor placed in front of the subject. A yellow circle (15-pixels in diameter) was an avatar for the research participant. Its position was determined by the joystick and was limited by the screen boundaries. A square shape (30 pixels in length) represented the prey. The movement of the prey was determined by a simple artificial intelligence algorithm (see below). Each trial ended with either the successful capture of the prey or after 20 sec, whichever came first. Successful capture was defined as any spatial overlap between the avatar circle and the prey square. Capture resulted in scored points; the number of points corresponded to prey color as follows: 1 point for orange; 3 points for green; 5 points for cyan.

The path of the prey was generated interactively using A-star pathfinding methods, which are commonly used in video gaming (Hart & Nils, 1968). For every frame (16.67 ms), we computed the cost of 15 possible future positions the prey could move to in the next time-step. These 15 positions were equally spaced on the circumference of a circle centered on the current position of the prey, with a radius equal to the maximum distance the prey could travel within one time-step. The cost in turn was based on the following two factors: the position in the field and the position of the avatar of the subject. The field that the prey moved in had a built-in bias for cost, which made the prey more likely to move toward the center. The cost due to distance from the avatar of the subject was transformed using a sigmoidal function: the cost became zero beyond a certain distance so that the prey did not move, and it became greater as distance from the avatar of the subject decreased. Eventually, the costs from these 15 positions were calculated and the position with the lowest cost was selected for the next movement. If the next movement was beyond the screen range (1,800 × 1,000 resolution), then the position with the second lowest cost was selected, and so on.

The maximum speed of the participant was 23 pixels per frame (and each frame was 16.67 ms). The maximum and minimum speeds of the prey remained the same across subjects. In trials with predators, a predator (triangle shape) appeared on 50% of trials. Capture by the predator led to points loss. Predators came in five different types (indicated by color) indicating different levels of points loss, ranging from 1 to 5 points.

The algorithm of the predator is to minimize the distance between itself and player. Unlike the prey, the predator algorithm is governed by this single rule. The design of the task reflects primarily the desire to have a rich and variegated virtual world with opportunities for choices at multiple levels that is neither trivially simple nor overly complex.

### Task presentation

Patients played at least 100 trials (average 119 trials) of the prey-pursuit task using a joystick (**Figure 1A, B**). The joystick was a commercially available joystick with a built-in potentiometer (Logitech Extreme Pro 3D). The joystick position was read out by a custom-coded program in Matlab running on the stimulus-control computer. The joystick was controlled by an algorithm that detected the positional change of the joystick and limited the maximum pixel movement to within 23 pixels in 16.67 ms. Task events were synchronized to the neural recording system via comments, sent through analog port, from the computer playing the task to the Neural Signal Processor (NSP) at 30 kHz.

### Linear–nonlinear model (LN-GLM)

To test the selectivity of neurons for various experimental variables, we constructed GLMs with navigational variables (Hardcastle et al., 2017; Pillow et al., 2008). The GLMs estimated the spike rate of one neuron during time bin *t* as an exponential function of the weighted sum of the relevant value of each variable at time *t*, for which the weights are determined by a set of coefficients (wi). The estimated firing rates from the GLMs can be expressed as follows:

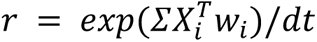

Where *r* denotes a vector of firing rates for one neuron over *T* time points across the session, and *i* indexes the variables of interest, for example, the position of the avatar on the screen. The vector of firing rates over *T* time points provide the benefit for modeling the neural activity without the need of specifically timelocking to a behavioral event. *Xi* is a matrix in which each column represents a set of ‘state variables’ of the subject obtained from binning the continuous variable so that all the columns for a particular row are 0, except for one column. In this case, state variables were obtained binning the position of self and prey over a 6x6 grid (36 bins). As a result, each neuron was assigned a design matrix with rows corresponding to the number of samples and columns corresponding to the number of bins.

Unlike conventional tuning curve analysis, GLM analysis does not assume the parametric shape of the tuning curve a priori. Instead, the weights, which define the shape of tuning for each neuron, were optimized by maximizing the Poisson log-likelihood of the observed spike train given the model-expected spike number, with additional regularization for the smoothness of parameters in a continuous variable, and a lasso regularization for parameters in a discrete variable. Position parameters were separately smoothed across rows and columns. The regularization hyperparameter was chosen by maximizing the cross-validation log-likelihood based on several randomly selected neurons. The unconstrained optimization with gradient and Hessian was performed (Matlab fminunc function). The model performance of each neuron was quantified by the log-likelihood of held out data under the model (**Figure 2C**). This cross-validation procedure was repeated ten times (tenfold cross-validation), and overfitting was penalized. Through multiple levels of penalties, we compared the performance of models with varying complexity.

### Forward model selection

Model selection was based on the cross-validated log-likelihood value for each model. We first fit *n* models with a single variable, where *n* is the total number of variables. The best single model was determined by the largest increase in spike-normalized log-likelihood from the null model (that is, the model with a single parameter representing the mean firing rate). Then, additional variables (*n – 1* in total) were added to the best single variable model. The best two variable model was preferred over the single variable model only if it significantly improved the cross-validation log-likelihood value (Wilcoxon signed-rank test, α = 0.05). Likewise, the procedure was continued for the three-variable model and beyond if adding more variables significantly improved the model performance, and the best, simplest model was selected. The cell was categorized as not tuned to any of the variables considered if the log-likelihood increase was not significantly higher than baseline, which was the mean firing rate of fitted neurons across the session.

### Spatial similarity index (SPAEF)

To compare the similarity between two spatial representations, we used the spatial efficiency measure (SPAEF) that prior literature suggests to be more robust than the 2D spatial correlation (Koch et al, 2018). It quantifies the similarity between two maps as follows:

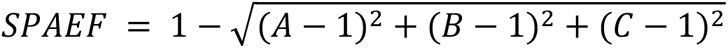

where A is the Pearson correlation between two maps, B is the ratio between the coefficients of variation for each map and C is the activity similarity measured by histogram profiles. Values near −1 indicate anticorrelated maps (one tends to be high when the other is low), 0 indicates uncorrelated maps and 1 indicates perfect matching between the two. By definition, SPAEF is not strictly constrained between -1 and 1. However, values outside this range tend to be rare, and in any case, in our data, values outside this interval never occurred.

To determine if the SPAEF values were significantly different from zero, we built a shuffled distribution over 1000 permutations. We then used a two-sided test to extract p-values, considering significance at p<0.05. To determine the noise ceiling, we performed 1000 within-agent half-splits. That is, for each iteration and each neuron, we randomly split the trials for each agent into two independent halves, computed the spatial tuning maps separately for each half, and then measured the spatial similarity between the two resulting maps. This process estimates the upper bound of explainable variance by accounting for the inherent noise in the data, providing a reference against which observed spatial similarities can be compared.

### K-means clustering and Principal Component Analysis

For this analysis, we included the entire neuronal population (390 neurons). We computed spatial tuning curves for each neuron based on the positions of the self and chosen prey. The task space (computer monitor) was divided into a 6×6 grid, resulting in a 36-bin tuning curve for each neuron per agent. Each tuning curve was then normalized between 0 and 1 on a per-neuron basis, by dividing by the maximum value of the respective neuron’s tuning curve. This ensured that differences in firing rates did not bias our subsequent analyses. We then applied the k-means clustering algorithm to the normalized tuning curves. Clustering was performed separately for self-position tuning curves and chosen prey-position tuning curves. To determine the optimal number of clusters we used silhouette values, which assess clustering quality by measuring how well each data point fits within its assigned cluster relative to other clusters. A silhouette score closer to -1 indicates potential misclassification, while a score closer to +1 suggests strong cluster cohesion and clear separation from other clusters (Rousseeuw, 1986). We selected the number of clusters that maximized the silhouette value, which turned out to be two for both self and chosen prey position tuning curves. Thus we could classify each neuron as belonging to cluster 1 or cluster 2.

We then performed principal component analysis (PCA) to visualize how the structure of the spatial tuning curves varies across neurons. That is, identifying the main patterns of variability in neurons’ responses to different positions. PCA was applied to the same normalized tuning curves (*neurons × bins* matrix) used for k-means clustering, where each row represented a neuron and each column corresponded to a spatial bin in its tuning curve. The resulting principal components (PCs) captured different aspects of tuning variability across the population.

To visualize this structure, we plotted neurons in the PCA space using their PC1 and PC2 scores and colored them according to their k-means cluster assignment. This allowed us to assess whether neurons with similar tuning properties grouped together in the PCA-defined space.

Specifically, we applied PCA separately to the self-position tuning maps and the chosen prey tuning maps, coloring neurons according to their respective k-means clustering results (**Figure 3C, D**). Lastly, we re-applied PCA to the chosen prey tuning maps and colored neurons by cross-referencing their cluster identities from both self and chosen prey tuning maps (**Figure 3E**). This final analysis highlighted differences in cluster membership between self and chosen prey tuning representations.

### Information per spike and SNR

This analysis was conducted on the raw (non-normalized) self-position spatial tuning curves (again 36 bins were used). Each neuron was assigned a cluster identity based on the results of k-means clustering applied to the self-position tuning curves. We calculated the spatial information per spike as the mutual information between the spatial location and the spike train, computed as:

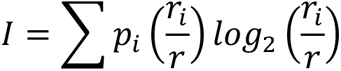

Where *r*i is the firing rate of each spatial bin (36 bins total) for neuron *n*, *r* is the average firing rate across all bins, and *p*i is the probability of occupying each spatial bin (Skaggs et al., 1992). In our case, since participants were able to sample the whole task space (**Figure 1B, C** and **Figure 3A**), we assumed uniform occupancy.

To assess the stability of cells firing, we calculate the signal-to-noise ratio (SNR), defined as the variance in the mean firing rate across bins divided by the mean rate itself. This metric provides a measure of the consistency of spatial encoding (Fenton and Muller, 1998).

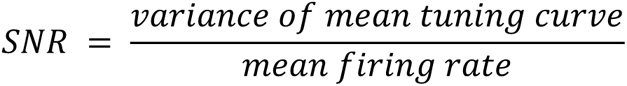

Higher SNR signifies consistency in firing across spatial locations, while lower SNR indicates less spatially structured responses. Significance of differences between groups was assessed using a non-parametric Wilcoxon rank-sum test.

### Spatial similarity matrix

To represent the correlation structure of self and prey representations, we constructed an N×N matrix for each agent, with N number of neurons. Each matrix entry represents the spatial similarity between the spatial maps of neuron pairs. Spatial maps were normalized between 0 and 1. The spatial similarity matrix is symmetric and contains values between -1 and +1.

For visualization purposes only, we applied hierarchical clustering to the self spatial similarity matrix. Specifically, we defined the dissimilarity self matrix as *1-spatial similarity matrix* and then applied the clustering algorithm using average linkage method. The extracted indices were then used to sort the matrices for the remaining agents (chosen and unchosen prey).

### Pairwise correlations

To compute the pairwise correlations between spatial similarity matrices, we used Pearson’s correlation coefficient. We assessed significance constructing a null distribution over 500 permutations. We then used a two-sided test to extract p-values, with significance considered at p<0.05

### Neural subspaces analysis

We performed PCA on the spatial similarity matrix by treating the self spatial similarity matrix as a covariance-like matrix and applying eigendecomposition. However, this matrix was not positive semi-definite (PSD). To approximate it to its nearest PSD form, we performed eigendecomposition and set negative eigenvalues to zero. We then identified the principal components (PCs) through a second eigendecomposition. Each identified self-PC represents an orthogonal direction in the N-dimensional neural space.

We then projected the prey spatial similarity matrices onto the self-PCs and quantified the percent of variance explained relative to the total variance of the self spatial similarity matrix (**Figure 4I, J**; green and grey bars). This procedure reveals the amount of self and prey variance shared in the self-PCs. Thus, if self and prey maps occupied semi-orthogonal subspaces, self-PCs would capture little variance from prey maps. Conversely, if the maps shared collinear subspaces, self-PCs would generalize across agents, explaining significant variance from prey representations.

To assess significance, we divided the neural data for each neuron into two halves (500 splits). For each split, spatial tuning curves were computed separately for each agent using one half of the data. We constructed the spatial similarity matrix, and identified the top self-PCs for the self training set (half 1). We then projected the self and prey maps onto the self PCs (halves 2 projected onto self half 1). The distribution of variance explained from within-self projections served as an estimate of noise ceiling. The variability of variance explained in within- and cross-condition projections (chosen prey and unchosen prey onto self-PCs) provides confidence intervals estimates. Variance explained in cross-condition projections was statistically compared to noise ceiling (assessed from the within-self variance distribution) using paired tests with Bonferroni correction.

Last, to quantify the degree of subspace orthogonality, we performed eigendecomposition on both the self and prey similarity matrices to identify the top self- and prey- PCs. We computed the angle between corresponding PCs using cosine similarity. Cosine similarity measures the cosine of the angle between two vectors, capturing their directional alignment in high-dimensional space. To determine statistical significance, we generated a null distribution via 500 permutations and computed permutation-based p-values using a two-tailed test, considering p < 0.05 as significant.

URL for Supplementary Video: https://www.youtube.com/watch?v=qR_54WB9XpI

## REFERENCES

Barak, O., Rigotti, M., & Fusi, S. (2013). The sparseness of mixed selectivity neurons controls the generalization–discrimination trade-off. Journal of Neuroscience, 33(9), 3844–3856.

Behrens, T. E., Muller, T. H., Whittington, J. C., Mark, S., Baram, A. B., Stachenfeld, K. L., & Kurth-Nelson, Z. (2018). What is a cognitive map? Organizing knowledge for flexible behavior. Neuron, 100(2), 490–509.

Bernardi, S., Benna, M. K., Rigotti, M., Munuera, J., Fusi, S., & Salzman, C. D. (2020). The geometry of abstraction in the hippocampus and prefrontal cortex. Cell, 183(4), 954–967.

Brown, T. I., Carr, V. A., LaRocque, K. F., Favila, S. E., Gordon, A. M., Bowles, B., … & Wagner, A. D. (2016). Prospective representation of navigational goals in the human hippocampus. Science, 352(6291), 1323–1326.

Burge, J., & Bonnen, K. (2025). Continuous psychophysics: past, present, future. Trends in Cognitive Sciences.

Cervera, R. L., Wang, M. Z., & Hayden, B. Y. (2020). Systems neuroscience of curiosity. Current Opinion in Behavioral Sciences, 35, 48–55.

Chaure, F. J., Rey, H. G. & Quian Quiroga, R. A novel and fully automatic spike-sorting implementation with variable number of features. J Neurophysiol 120, 1859–1871 (2018).

Chersi, F., & Burgess, N. (2015). The cognitive architecture of spatial navigation: hippocampal and striatal contributions. Neuron, 88(1), 64–77.

Cisek, P., & Kalaska, J. F. (2010). Neural mechanisms for interacting with a world full of action choices. Annual review of neuroscience, 33(1), 269–298.

Dale, A.M., Fischl, B., Sereno, M.I., 1999. Cortical surface-based analysis. I. Segmentation and surface reconstruction. Neuroimage 9, 179–194.

Dalton, M. A., D’Souza, A., Lv, J., & Calamante, F. (2022). New insights into anatomical connectivity along the anterior–posterior axis of the human hippocampus using in vivo quantitative fibre tracking. elife, 11, e76143.

Danjo, T., Toyoizumi, T., & Fujisawa, S. (2018). Spatial representations of self and other in the hippocampus. Science, 359(6372), 213–218.

Diehl, G. W., Hon, O. J., Leutgeb, S., & Leutgeb, J. K. (2017). Grid and nongrid cells in medial entorhinal cortex represent spatial location and environmental features with complementary coding schemes. Neuron, 94(1), 83–92.

Ebitz, R. B., & Hayden, B. Y. (2021). The population doctrine in cognitive neuroscience. Neuron, 109(19), 3055–3068.

Ebitz, R. B., Sleezer, B. J., Jedema, H. P., Bradberry, C. W., & Hayden, B. Y. (2019). Tonic exploration governs both flexibility and lapses. PLoS computational biology, 15(11), e1007475.

Eisenreich, B. R., Hayden, B. Y., & Zimmermann, J. (2019). Macaques are risk-averse in a freely moving foraging task. Scientific reports, 9(1), 15091.

Ekstrom, A. D., Kahana, M. J., Caplan, J. B., Fields, T. A., Isham, E. A., Newman, E. L., & Fried, I. (2003). Cellular networks underlying human spatial navigation. Nature, 425(6954), 184–188.

Elsayed, G. F., Lara, A. H., Kaufman, M. T., Churchland, M. M., & Cunningham, J. P. (2016). Reorganization between preparatory and movement population responses in motor cortex. Nature communications, 7(1), 13239.

Epstein, R. A., Patai, E. Z., Julian, J. B., & Spiers, H. J. (2017). The cognitive map in humans: spatial navigation and beyond. Nature neuroscience, 20(11), 1504–1513.

Fabian, S. T., Sumner, M. E., Wardill, T. J., Rossoni, S., & Gonzalez-Bellido, P. T. (2018). Interception by two predatory fly species is explained by a proportional navigation feedback controller. Journal of The Royal Society Interface, 15(147), 20180466.

Fenton, A. A., & Muller, R. U. (1998). Place cell discharge is extremely variable during individual passes of the rat through the firing field. Proceedings of the National Academy of Sciences, 95(6), 3182–3187.

Fine, J. M., Maisson, D. J. N., Yoo, S. B. M., Cash-Padgett, T. V., Wang, M. Z., Zimmermann, J., & Hayden, B. Y. (2023). Abstract value encoding in neural populations but not single neurons. Journal of Neuroscience, 43(25), 4650–4663.

Fine, J. M., Yoo, S. B. M., & Hayden, B. Y. (2024). Control over a mixture of policies determines change of mind topology during continuous choice. bioRxiv, 2024-04.

Forli, A., & Yartsev, M. M. (2023). Hippocampal representation during collective spatial behaviour in bats. Nature, 621(7980), 796–803. Freesurfer: https://freesurfer.net/fswiki/FreeSurferMethodsCitation

Fusi, S., Miller, E. K., & Rigotti, M. (2016). Why neurons mix: high dimensionality for higher cognition. Current opinion in neurobiology, 37, 66–74.

Gauthier, J. L., & Tank, D. W. (2018). A dedicated population for reward coding in the hippocampus. Neuron, 99(1), 179–193.

Gordon, J., Maselli, A., Lancia, G. L., Thiery, T., Cisek, P., & Pezzulo, G. (2021). The road towards understanding embodied decisions. Neuroscience & Biobehavioral Reviews, 131, 722–736.

Groppe DM, Bickel S, Dykstra AR, et al. iELVis: An open source MATLAB toolbox for localizing and visualizing human intracranial electrode data. J Neurosci Methods. 2017;281:40–48.

Hardcastle, K., Maheswaranathan, N., Ganguli, S., & Giocomo, L. M. (2017). A multiplexed, heterogeneous, and adaptive code for navigation in medial entorhinal cortex. Neuron, 94(2), 375–387.

Hart, P. E. & Nils, J. Formal basis for the heuristic determination of minumum cost path. IEEE Trans. Syst. Sci. Cyber. 4, 100–107 (1968).

Harvey, C. D., Collman, F., Dombeck, D. A., & Tank, D. W. (2009). Intracellular dynamics of hippocampal place cells during virtual navigation. Nature, 461(7266), 941–946.

Hayden, B. Y. (2019). Why has evolution not selected for perfect self-control?. Philosophical Transactions of the Royal Society B, 374(1766), 20180139.

Jacobs, Joshua, Michael J. Kahana, Arne D. Ekstrom, Matthew V. Mollison, and Itzhak Fried. "A sense of direction in human entorhinal cortex." Proceedings of the National Academy of Sciences 107, no. 14 (2010): 6487–6492.

Jenkinson, M., Bannister, P., Brady, J. M. and Smith, S. M. 2002. Improved Optimisation for the Robust and Accurate Linear Registration and Motion Correction of Brain Images. NeuroImage, 17(2), 825–841.

Jenkinson, M., & Smith, S. (2001). A global optimisation method for robust affine registration of brain images. Medical image analysis, 5(2), 143–156.

Johnston, W. J., Fine, J. M., Yoo, S. B. M., Ebitz, R. B., & Hayden, B. Y. (2024). Semi-orthogonal subspaces for value mediate a binding and generalization trade-off. Nature Neuroscience, 27(11), 2218–2230.

Joshi, A., Scheinost, D., Okuda, H., Belhachemi, D., Murphy, I., Staib, L. H., & Papademetris, X. (2011). Unified framework for development, deployment and robust testing of neuroimaging algorithms. Neuroinformatics, 9(1), 69–84.

Jung, M. W., Wiener, S. I., & McNaughton, B. L. (1994). Comparison of spatial firing characteristics of units in dorsal and ventral hippocampus of the rat. Journal of Neuroscience, 14(12), 7347–7356.

Kaufman, M. T., Benna, M. K., Rigotti, M., Stefanini, F., Fusi, S., & Churchland, A. K. (2022). The implications of categorical and category-free mixed selectivity on representational geometries. Current opinion in neurobiology, 77, 102644.

Kjelstrup, K. B., Solstad, T., Brun, V. H., Hafting, T., Leutgeb, S., Witter, M. P., … & Moser, M. B. (2008). Finite scale of spatial representation in the hippocampus. Science, 321(5885), 140–143.

Koch, J., Demirel, M. C., & Stisen, S. (2018). The SPAtial EFficiency metric (SPAEF): Multiple-component evaluation of spatial patterns for optimization of hydrological models. Geoscientific Model Development, 11(5), 1873–1886.

Kunz, L., Brandt, A., Reinacher, P. C., Staresina, B. P., Reifenstein, E. T., Weidemann, C. T., … & Jacobs, J. (2021). A neural code for egocentric spatial maps in the human medial temporal lobe. Neuron, 109(17), 2781–2796.

Li X, Morgan PS, Ashburner J, Smith J, Rorden C (2016) The first step for neuroimaging data analysis: DICOM to NIfTI conversion. J Neurosci Methods. 264:47–56. doi: 10.1016/j.jneumeth.2016.03.001. PMID: 26945974

Mackay, S., Reber, T. P., Bausch, M., Boström, J., Elger, C. E., & Mormann, F. (2024). Concept and location neurons in the human brain provide the ‘what’and ‘where’in memory formation. Nature Communications, 15(1), 7926.

Magnotti JF, Wang Z, Beauchamp MS. RAVE: comprehensive open-source software for reproducible analysis and visualization of intracranial EEG data. NeuroImage (2020) 223:117341.

Maguire, E. A., Woollett, K., & Spiers, H. J. (2006). London taxi drivers and bus drivers: a structural MRI and neuropsychological analysis. Hippocampus, 16(12), 1091–1101.

Martinez-Trujillo, J. (2025). Why do primates have view cells instead of place cells?. Trends in Cognitive Sciences.

Merel, J., Botvinick, M., & Wayne, G. (2019). Hierarchical motor control in mammals and machines. Nature communications, 10(1), 5489.

Miller, J. F., Neufang, M., Solway, A., Brandt, A., Trippel, M., Mader, I., … & Schulze-Bonhage, A. (2013). Neural activity in human hippocampal formation reveals the spatial context of retrieved memories. Science, 342(6162), 1111–1114.

Nyberg, N., Duvelle, É., Barry, C., & Spiers, H. J. (2022). Spatial goal coding in the hippocampal formation. Neuron, 110(3), 394–422.

O’Keefe, J., Burgess, N., Donnett, J. G., Jeffery, K. J., & Maguire, E. A. (1998). Place cells, navigational accuracy, and the human hippocampus. Philosophical Transactions of the Royal Society of London.

O’Keefe, J., & Dostrovsky, J. (1971). The hippocampus as a spatial map: preliminary evidence from unit activity in the freely-moving rat. Brain research.

Olberg, R. M., Worthington, A. H., & Venator, K. R. (2000). Prey pursuit and interception in dragonflies. Journal of Comparative Physiology A, 186, 155–162.

Omer, D. B., Maimon, S. R., Las, L., & Ulanovsky, N. (2018). Social place-cells in the bat hippocampus. Science, 359(6372), 218–224.

Panichello, M. F., & Buschman, T. J. (2021). Shared mechanisms underlie the control of working memory and attention. Nature, 592(7855), 601–605.

Parthasarathy, A., Herikstad, R., Bong, J. H., Medina, F. S., Libedinsky, C., & Yen, S. C. (2017). Mixed selectivity morphs population codes in prefrontal cortex. Nature neuroscience, 20(12), 1770–1779.

Pillow, J. W. et al. Spatio-temporal correlations and visual signalling in a complete neuronal population. Nature 454, 995–999 (2008).

Poppenk, J., Evensmoen, H. R., Moscovitch, M., & Nadel, L. (2013). Long-axis specialization of the human hippocampus. Trends in cognitive sciences, 17(5), 230–240.

Poucet, B., & Hok, V. (2017). Remembering goal locations. Current opinion in behavioral sciences, 17, 51–56.

Rao, R. P., von Heimendahl, M., Bahr, V., & Brecht, M. (2019). Neuronal responses to conspecifics in the ventral CA1. Cell reports, 27(12), 3460–3472.

Forli, A., & Yartsev, M. M. (2023). Hippocampal representation during collective spatial behaviour in bats. Nature, 621(7980), 796–803.

Rigotti, M., Barak, O., Warden, M. R., Wang, X. J., Daw, N. D., Miller, E. K., & Fusi, S. (2013). The importance of mixed selectivity in complex cognitive tasks. Nature, 497(7451), 585–590.

Rousseeuw, P. J. (1987). Silhouettes: a graphical aid to the interpretation and validation of cluster analysis. Journal of computational and applied mathematics, 20, 53–65.

Skaggs, W., Mcnaughton, B., & Gothard, K. (1992). An information-theoretic approach to deciphering the hippocampal code. Advances in neural information processing systems, 5.

Stangl, M., Topalovic, U., Inman, C. S., Hiller, S., Villaroman, D., Aghajan, Z. M., … & Suthana, N. (2021). Boundary-anchored neural mechanisms of location-encoding for self and others. Nature, 589(7842), 420–425.

Stephens, D. W., & Krebs, J. R. (1986). Foraging theory (Vol. 6). Princeton university press.

Strange, B. A., Witter, M. P., Lein, E. S., & Moser, E. I. (2014). Functional organization of the hippocampal longitudinal axis. Nature reviews neuroscience, 15(10), 655–669.

Suthana, N. A., Ekstrom, A. D., Moshirvaziri, S., Knowlton, B., & Bookheimer, S. Y. (2009). Human hippocampal CA1 involvement during allocentric encoding of spatial information. Journal of Neuroscience, 29(34), 10512–10519.

Tang, C., Herikstad, R., Parthasarathy, A., Libedinsky, C., & Yen, S. C. (2020). Minimally dependent activity subspaces for working memory and motor preparation in the lateral prefrontal cortex. Elife, 9, e58154.

To, T. V., Wang, D. X., Wolfe, C. B., & Lega, B. C. (2024). Neurophysiological Evidence of Human Hippocampal Longitudinal Differentiation in Associative Memory.

Tye, K. M., Miller, E. K., Taschbach, F. H., Benna, M. K., Rigotti, M., & Fusi, S. (2024). Mixed selectivity: Cellular computations for complexity. Neuron, 112(14), 2289–2303.

Wang, M. Z., & Hayden, B. Y. (2021). Latent learning, cognitive maps, and curiosity. Current Opinion in Behavioral Sciences, 38, 1–7.

Watrous, A. J., Miller, J., Qasim, S. E., Fried, I., & Jacobs, J. (2018). Phase-tuned neuronal firing encodes human contextual representations for navigational goals. Elife, 7, e32554.

Wilson, M. A., & McNaughton, B. L. (1993). Dynamics of the hippocampal ensemble code for space. Science, 261(5124), 1055–1058.

Xie, Y., Hu, P., Li, J., Chen, J., Song, W., Wang, X. J., … & Wang, L. (2022). Geometry of sequence working memory in macaque prefrontal cortex. Science, 375(6581), 632–639.

Yacoub, E., Grier, M. D., Auerbach, E. J., Lagore, R. L., Harel, N., Adriany, G., … & Zimmermann, J. (2020). Ultra-high field (10.5 T) resting state fMRI in the macaque. Neuroimage, 223, 117349.

Yang, A. I., Wang, X., Doyle, W. K., Halgren, E., Carlson, C., Belcher, T. L., et al. (2012). Localization of dense intracranial electrode arrays using magnetic resonance imaging. NeuroImage 63(1), 157–165. doi:10.1016/j.neuroimage.2012.06.039

Ydenberg, R. C., & Dill, L. M. (1986). The economics of fleeing from predators. In Advances in the Study of Behavior (Vol. 16, pp. 229-249). Academic Press.

Yoo, S. B. M., Tu, J. C., Piantadosi, S. T., & Hayden, B. Y. (2020). The neural basis of predictive pursuit. Nature neuroscience, 23(2), 252–259.

Yoo, S. B. M., Hayden, B. Y., & Pearson, J. M. (2021A). Continuous decisions. Philosophical Transactions of the Royal Society B, 376(1819), 20190664.

Yoo, S. B. M., Tu, J. C., & Hayden, B. Y. (2021B). Multicentric tracking of multiple agents by anterior cingulate cortex during pursuit and evasion. Nature communications, 12(1), 1985.

Yoo, S. B. M., & Hayden, B. Y. (2020). The transition from evaluation to selection involves neural subspace reorganization in core reward regions. Neuron, 105(4), 712–724.

Zhang, X., Cao, Q., Gao, K., Chen, C., Cheng, S., Li, A., … & Miao, C. (2024). Multiplexed representation of others in the hippocampal CA1 subfield of female mice. Nature Communications, 15(1), 3702.

